# Direct visualization of Na,K-ATPase clustering by 3D DNA-PAINT MINFLUX nanoscopy

**DOI:** 10.64898/2026.06.30.735534

**Authors:** Bruno Stojcic, Ana Agostinho, Luca Panconi, Hans Blom, Hjalmar Brismar

## Abstract

Direct validation of the nanoscale structural organization of membrane proteins requires localization precision that matches their molecular dimensions. The sodium-potassium pump, or the Na,K-ATPase is an integral membrane protein responsible for maintaining electrochemical gradients and cellular energy homeostasis. Although its crystal structure is characterized, the organization of the Na,K-ATPase within native plasma membranes, particularly whether it forms functional oligomers, remains an open question. Here, we combined 3D MINFLUX nanoscopy with DNA-PAINT with sub-10 nm localization precision to map the clustering topology of the Na,K-ATPase in mammalian cells. By targeting EGFP-tagged Na,K-ATPase α1 and β1 subunits using anti-GFP nanobodies, we obtained high-density 3D localization maps of the protein in the plasma membrane. To evaluate the point patterns, we developed a computational data-driven spatial point assignment approach that segments apical and basal localizations, mitigating clustering artifacts produced by imaging two membranes in close proximity. Furthermore, we used a spatial statistical approach analyzing sequential nearest-neighbour distances to elucidate supramolecular arrangement information. Our data reveal a preferential nearest-neighbour distance of approximately 7 nm, providing direct visual confirmation of Na,K-ATPase dimerization. Additionally, we identified higher-order nanoclusters composed of up to 21 proteins. These findings provide definitive structural evidence of the dimeric configuration of Na,K-ATPase, establishing a foundation for future research on the functional and regulatory implications of Na,K-ATPase clustering.

## Introduction

The Na,K-ATPase is an integral plasma membrane protein expressed in all eukaryotic cells. Na,K-ATPase is a key regulator of the membrane potential by exporting three Na^+^ and importing two K^+^ ions per hydrolyzed ATP molecule.^1^ This function accounts for 30% to 70% of cellular energy expenditure. Functional Na,K-ATPase is a heterodimer consisting of the catalytic alpha (*α*) subunit, a glycosylated regulatory beta subunit (*β*), and an auxiliary gamma (*γ*) subunit. Both alpha and beta subunits have four isoforms (*α*1-4 and *β*1-4), with tissue specific expression patterns.^2^ The *α*1 and *β*1 isoforms are ubiquitously expressed and predominate in kidney epithelia.^3^

Despite characterization of Na,K-ATPase structure and function since its discovery in 1957, the nanoscale spatial organisation of the enzyme in the plasma membrane and its functional implications remain under-investigated.^4^ Na,K-ATPase self-interaction and functional dimer formation have been proposed and debated.^3^ Experimental evidence for self-interaction of Na,K-ATPase remains indirect. Using Förster Resonance Energy Transfer combined with Fluorescence Correlation Spectroscopy (FRET-FCS) in living cells, we demonstrated that 34% +/−11% of Na,K-ATPase exists in a dimer configuration.^5^ Furthermore, Single Molecule Localization Microscopy (PALM/STORM) studies have provided insight into the higher-level plasma membrane distribution of the Na,K-ATPase.^6,7^

Methods such as FRET-FCS and camera-based PALM/STORM either provide indirect evidence based on fluorophore proximity or diffusion dynamics, or do not achieve sufficient resolution to elucidate the structural distribution of Na,K-ATPase. Visualizing Na,K-ATPase at a single-molecule scale in its native membrane environment has not yet been achieved.

To overcome the diffraction limit and resolve Na,K-ATPase architecture in the native cell membrane, we employ super-resolution microscopy, specifically, Minimal Photon Fluxes (MINFLUX) nanoscopy. MINFLUX offers 3D spatial localisation precision below 10 nm.^8,9^ We combined MINFLUX acquisition with DNA-PAINT (DNA-based Point Accumulation for Imaging in Nanoscale Topography).^10^ DNA-PAINT enables repeated acquisition of high-density localization maps, a requirement for resolving packed molecules within nanoscale clusters. While 3D MINFLUX provides the necessary sub-10 nm localization precision, it relies on the dense, non-photobleaching target sampling afforded by DNA-PAINT to reliably resolve adjacent proteins without undercounting or spatial overlap. Expressing Na,K-ATPase isoforms as an EGFP fusion protein and labeling it with an anti-GFP nanobody conjugated to a DNA oligonucleotide (docking strand) allows MINFLUX acquisition by imaging cells in a solution containing complementary DNA strands coupled to fluorophores (imager strands). This approach enables repeated localizations, provides the longer fluorophore on-times required for MINFLUX localizations, and overcomes photobleaching limitations.^11^

Here, we apply 3D MINFLUX nanoscopy combined with DNA-PAINT on A498, a human kidney carcinoma cell line transiently expressing either *α*1 or *β*1 Na,K-ATPase sub-units N-terminally tagged with EGFP. We provide direct, quantitative visualization of Na,K-ATPase spatial organization to confirm the existence and configuration of Na,K-ATPase dimers and nanoscale clusters. To analyze the resulting high-resolution point cloud data and quantify Na,K-ATPase clustering, we employ spatial statistics, specifically nearest-neighbour analysis. We introduce an extension of nearest-neighbour analysis to systematically study sequential distance changes from the first to the n-th nearest-neighbour. This permits unbiased cluster analysis without prior knowledge of target structure and distribution. Finally, we develop and employ a data-driven point cloud separation algorithm to classify localizations to either the apical or basal membrane. This mitigates clustering artifacts associated with imaging membranes in close proximity, providing a tool applicable to general 3D point pattern data for membrane classification or investigating per-membrane clustering differences.

## Results

To perform DNA-PAINT MINFLUX, we expressed either Na,K-ATPase *α*1-EGFP or Na,K-ATPase *β*1-EGFP in A498 cells and immunostained the target protein using an anti-GFP nanobody conjugated to a DNA docking strand (*Figure 1 a*). Based on confocal imaging of the GFP signal, cells exhibiting wild type morphology and homogenous expression were selected for nanoscopy (*Figure 1 b, Supplementary Figure 1*). A confocal reference scan on the MINFLUX instrument was acquired to select regions with appropriate stability, utilizing in-focus gold beads close to the coverslip to ensure imaging of a thin membrane at the edge of the cell. Then, a 3×3 µm area was selected for a 3D MINFLUX acquisition (*Figure 1 c*). Pooled analysis of all acquisitions gave a median localization precision (σ) of σ_x_ = 4.64 nm, σ_y_ = 4.76 nm, and σ_z_ = 2.81 nm (*Figure 1 d*). Based on these data, a conservative 3 nm search radius threshold was established for trace ID reassignment before further analysis, in addition to a minimum distance threshold between adjacent points for all simulated point patterns.

**Figure 1.**
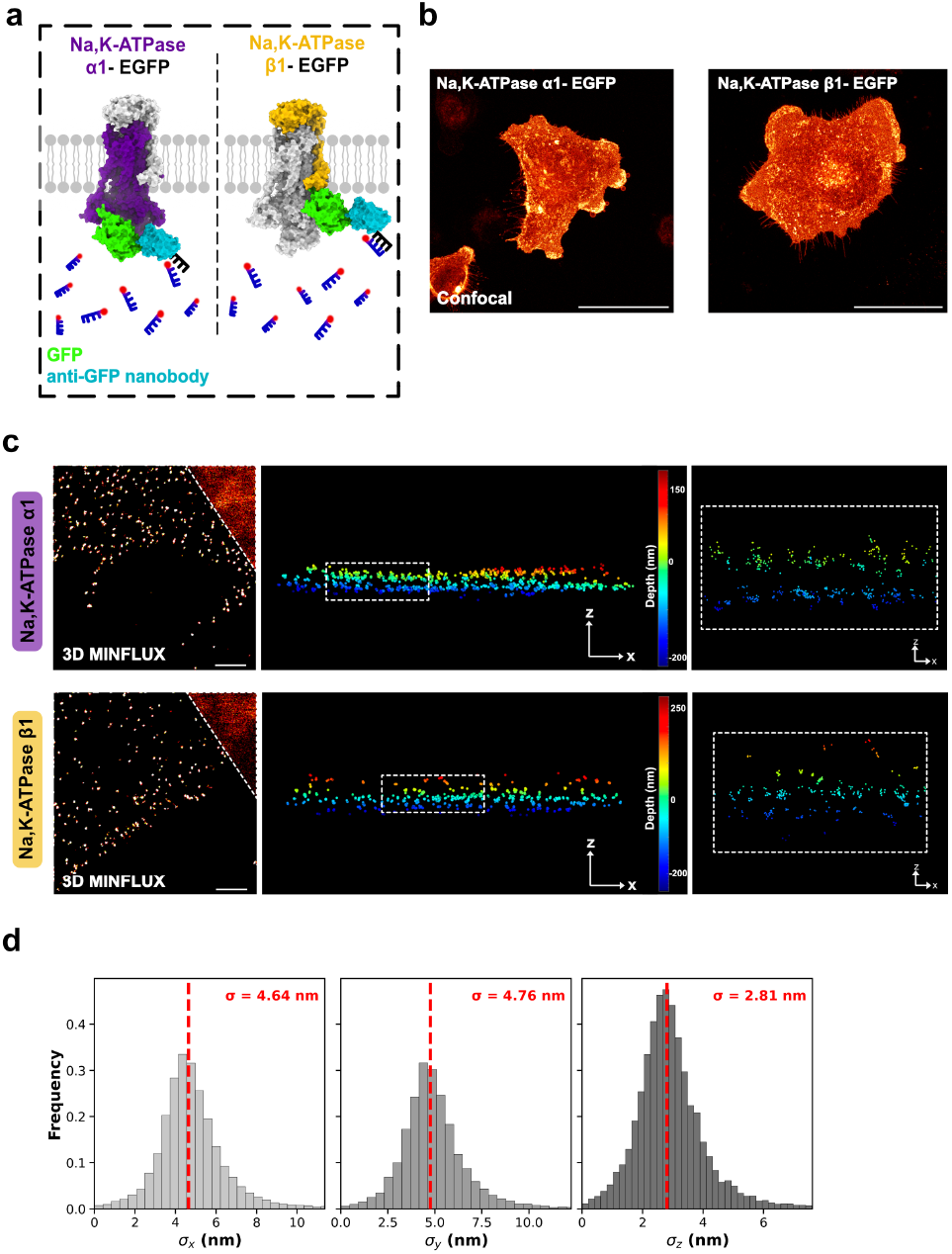
3D DNA-PAINT MINFLUX imaging of Na,K ATPase subunits. a. Schematic of labelling strategy used to perform DNA-PAINT based MINFLUX detection of *α*1 (purple) and *β*1 (yellow) subunits. Subunits are N-terminally tagged with EGFP (green). The tag is detected with an anti-GFP nanobody (cyan) coupled to a docking strand (black) and imaged using complementary imager strands (blue) conjugated to Abberior 660 dye b. Maximum intensity projections of cells expressing either *α*1-EGFP or *β*1-EGFP 72-hours post transfection, showing uniform membrane expression (scale bar: 50 µm). c. Confocal images used for cell selection and localization of a 3×3 µm region of interest. Spatial localisations are rendered in lateral (x,y) and axial (x,z) views (scale bar: 500 nm). 3D MINFLUX resolves proteins on opposing membranes spaced approximately 100 nm apart. d. Localisation precision distribution with a median σ of 4.64 nm in x, 4.76 nm in y, and 2.81 nm in z. A conservative 3 nm threshold defines the search radius for trace id reassignment.

The localizations were filtered based on trace length, effective frequency (efo) and center frequency ratio (cfr), and trace ID reassignment was performed via DBSCAN. Finally, three independent 1×1 µm regions of interest (ROIs) were selected per recording for downstream analysis (*Figure 2 a*). We calculated the first nearest-neighbour distance (NND) for each point in the selected ROIs, both the *α*1 and *β*1 subunits showed a preferential NND of approximately 7 nm (*Figure 2 b*). Quantification showed that 70% of points for *α*1 and 72% of points for *β*1 localized within a 10 nm radius of the first nearest-neighbour, confirming that this preferential distance is a consistent feature across datasets (*Figure 2 c*). To evaluate spatial randomness, the median observed number of points per ROI was used to simulate complete spatial randomness (CSR) in a 1×1 µm space, incorporating the observed z-distribution to model the third dimension (*Figure 2 d*). CSR was simulated a 1000 times and the resulting median distances (68 nm for *α*1and 75 nm for *β*1) were compared to the observed values, confirming that the observed preferential distances reflect non-random molecular organisation. (*Figure 2 e, Supplementary Figure 2*).

**Figure 2.**
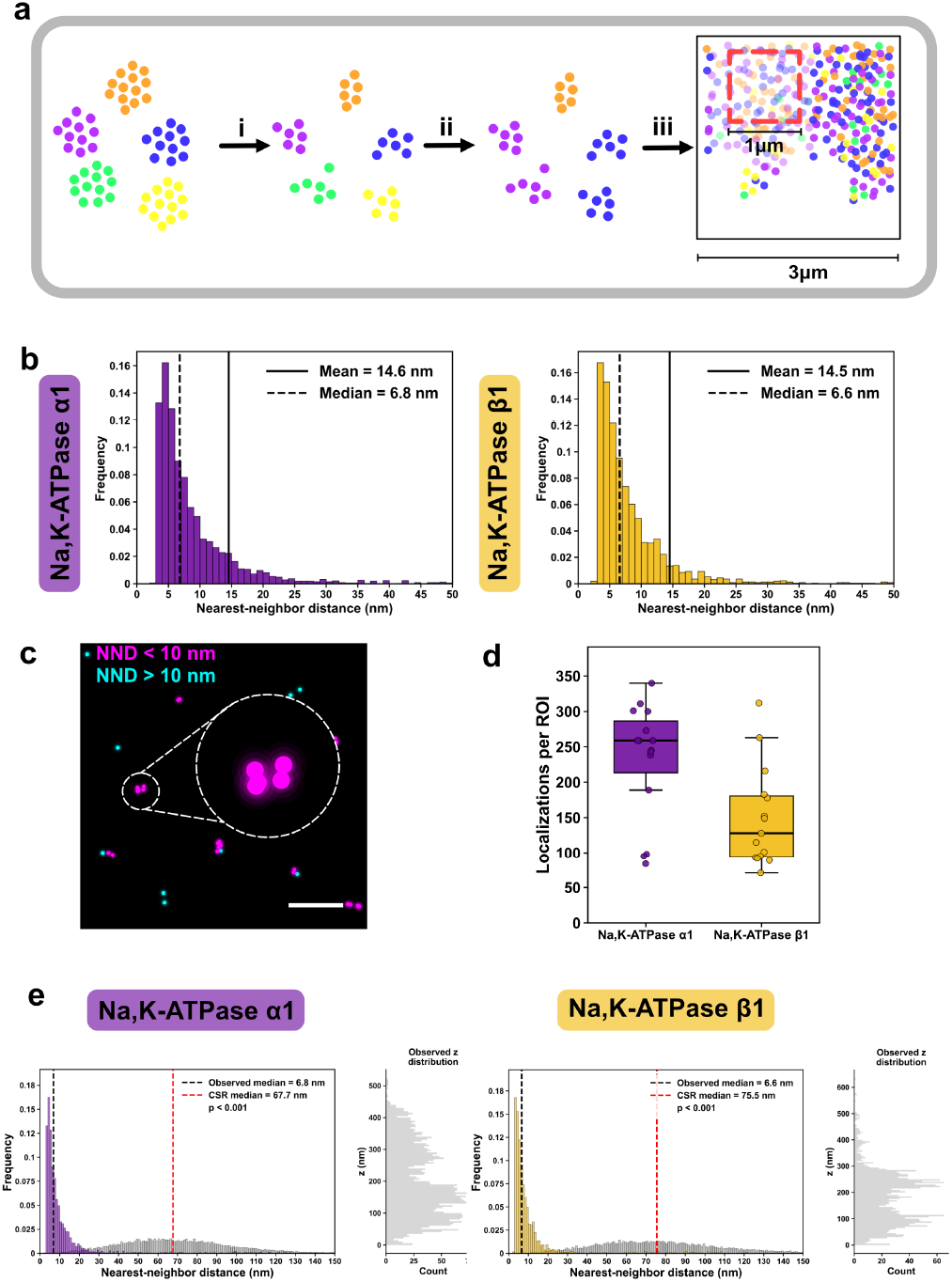
3D MINFLUX Nearest-Neighbour Analysis indicating a 7 nm preferential distance. **a**. Data processing workflow showing (i) filtering of raw localizations by effective frequency at offset (efo), center frequency ratio (cfr), and trace length. (ii) trace id reassignment via DBSCAN (min pts= 1, eps= 3 nm) to mitigate reblinking artifacts, and (iii) isolation of 1×1 µm regions of interest. **b**. First nearest-neighbour distance distributions showing a preferential separation of approximately 7 nm for both *α*1 and *β*1 subunits. **c**. Representative region of interest highlighting localizations with a nearest-neighbour distance under 10 nm (magenta) or over 10 nm (cyan) (scale bar: 100 nm). Closely spaced points indicate a homodimer configuration. **d**. Distribution of the number of localizations per region of interest used to define parameters for complete spatial randomness (CSR) simulations. **e**. Axial distribution of points applied to the three-dimensional simulations. Comparison of observed and simulated density curves demonstrates that the experimental distributions are non-random (p < 0.001).

To isolate per-membrane clustering patterns and mitigate artifacts arising from the close proximity of opposing plasma membranes, we implemented a data-driven manifold learning approach. This algorithm extracts the membrane topography from the input data and fits representative manifolds to the 3D point cloud (*Figure 3 a*), allowing each localisation to be classified as apical, basal, or an outlier (*Figure 3 b*). We computed first nearest-neighbour distances by treating the two membranes as separate datasets and compared the distributions to 2D CSR simulations. Although no pronounced per-membrane differences were observed, the median first NNDs were slightly longer (0.1-0.5 nm) compared to the previous unseparated analysis. This indicates that unseparated 3D data contained artificial trans-membrane nearest-neighbours. These results confirm the organisational trend observed in the unseparated 3D data, for *α*1, 64% (apical) and 71 % (basal) of points localize within a 10 nm radius of their first nearest-neighbour. Similarly, for *β*1, 69% (apical) and 76% (basal) of points have a first nearest-neighbour within this 10 nm radius. The median first nearest-neighbour distances differed significantly from the corresponding CSR simulations, confirming non-random preferential distributions at each membrane surface (*Figure 3 c, Supplementary Figure 3*).

**Figure 3.**
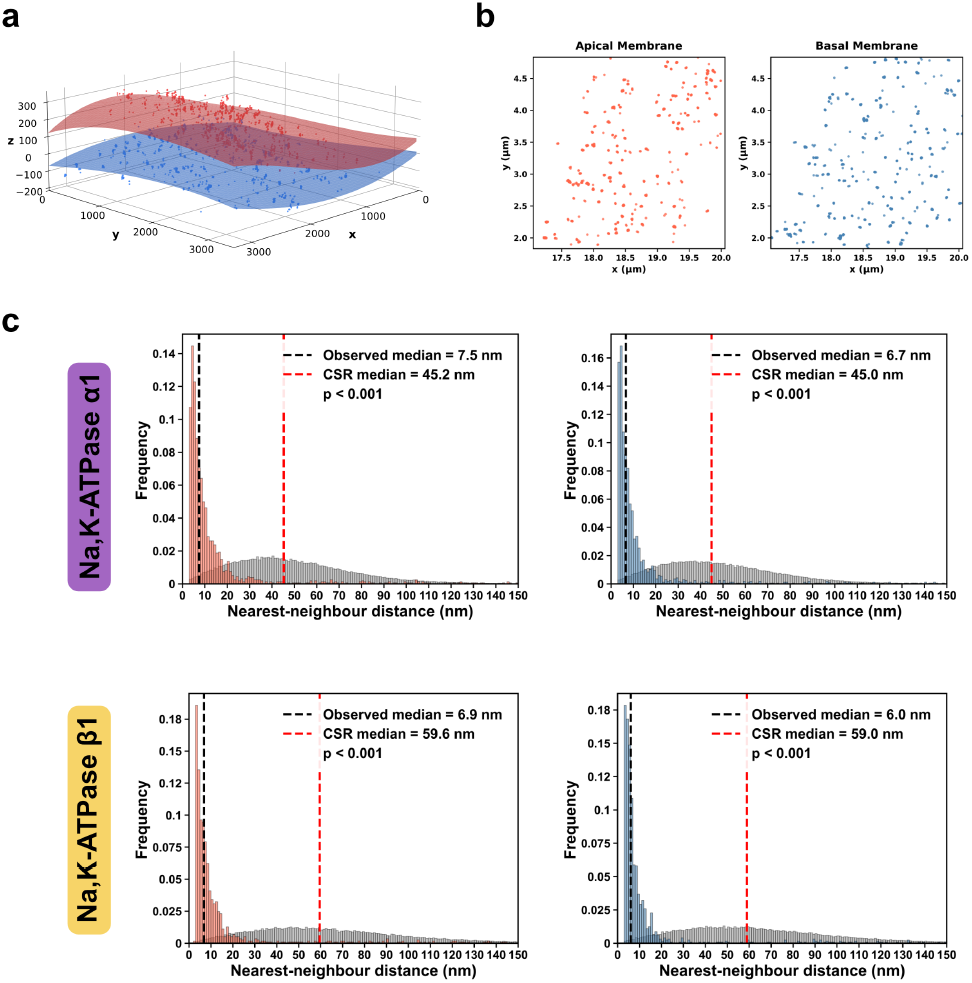
Membrane-specific nearest-neighbour analysis via data-driven manifold learning. a. Three-dimensional coordinates fitted with distinct apical and basal mathematical surfaces. b. Classification of individual localizations into apical or basal membrane datasets for independent spatial analysis. c. First nearest-neighbour distance distributions for each membrane surface compared against 1000 complete spatial randomness (CSR) simulations. Both subunits show a preferential distance between 6 and 7 nm on both surfaces (p< 0.001). Separation reveals a minor increase in the median distance, indicating the elimination of trans-membrane coordinate artifacts present in unseparated data.

We then extended the analysis to include the first 10 nearest neighbours for membrane-separated *α*1 and *β*1, comparing the results to the 2D CSR simulations (*Figure 4 a*). Since no statistical difference in the clustering was resolved between the two isoforms, we continued to probe the higher-order clustering of the *α*1 subunit, which showed a deviation in the NND-k trend curve compared to the CSR. The *α*1 NND curve showed an inflection point characterized by an increase in the nearest-neighbour distance between the fourth and the fifth neighbour generations (*Figure 4 b*). We investigated this behavior by developing an algorithm that simulates the spatial distribution of points in a 1×1 µm 2D space. The algorithm takes three inputs: observed median point density, observed median first NND, and labeling efficiency. It identifies the best fitting model relative to the observed data by testing varying ratios of dispersed versus clustered points (0-100%), evaluating a range of cluster sizes (2-n) assuming a normal distribution, adjusting the variance (σ) of the gaussian spatial distribution to match the experimental median first NND, and accounting for labeling efficiency by extrapolating the expected true number of points via the product of the observed density and the inverse of the labeling efficiency. Models that minimize the sum of squared errors (SSE) were selected, providing predictions for maximum cluster size, average cluster size, and the degree of clustering (*Figure 4 c*). The models generated improved fits when the corrected labeling density was applied, indicated by a significant overall reduction in the range of SSE values *(Figure 4 d-e*). The best fitting models for Na,K-ATPase *α*1 indicate maximum cluster sizes between 13 and 21 points, with high degree of clustering, between 55-100% (*Supplementary Table 2-3*). This demonstrates that traditional density-based methods, such as DBSCAN, which omit corrections for undetected molecules, underestimate cluster sizes and reflect the distribution of the detected points rather than the underlying biological architecture. Accounting for incomplete labelling is therefore essential when evaluating the nanoscale stoichiometry of uncharacterized membrane targets such as the Na,K-ATPase.

**Figure 4.**
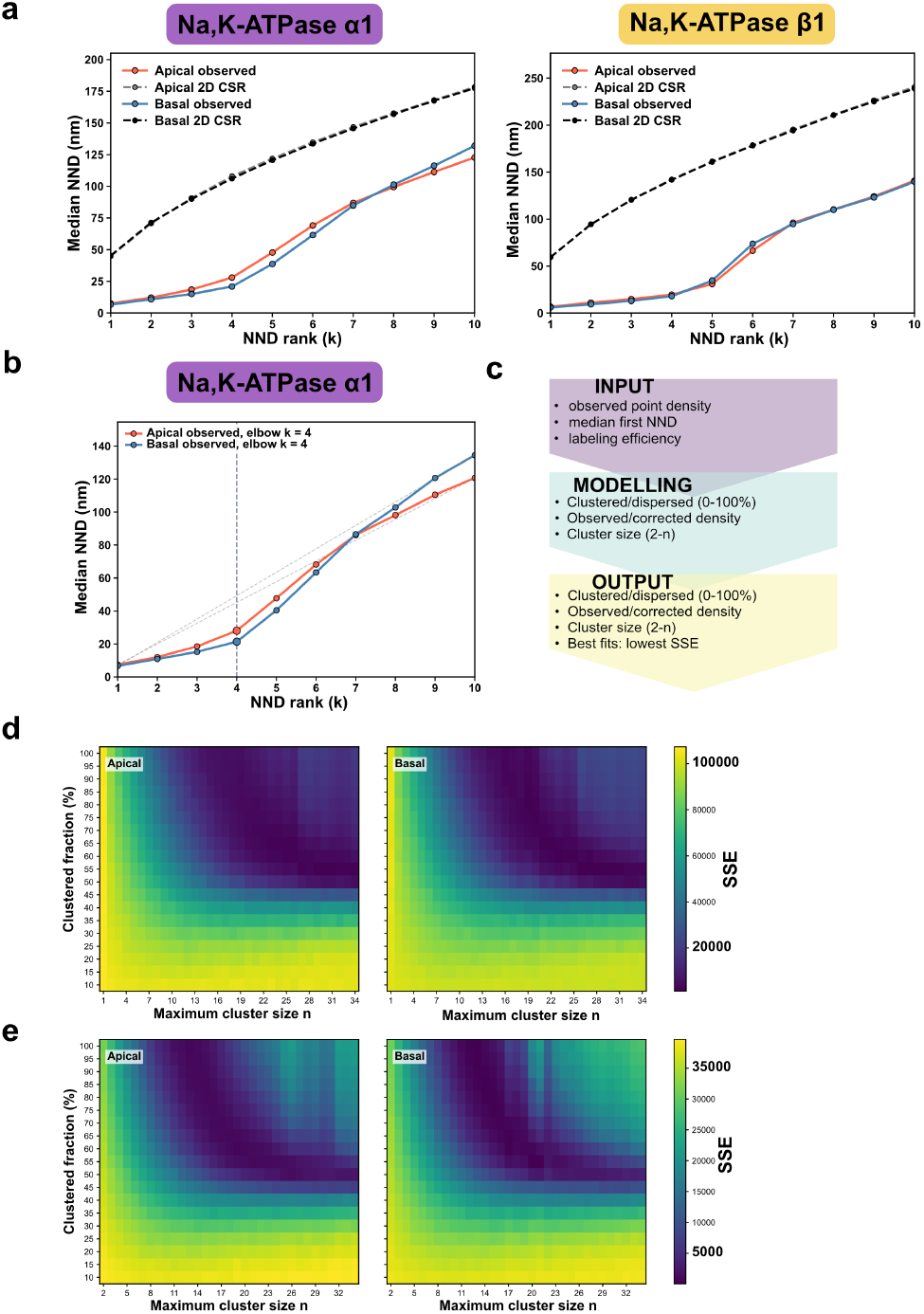
Nearest-neighbour analysis of higher-order nanocluster sizes. **a**. Median k-th nearest-neighbour distances for membrane-separated *α*1 and *β*1 subunits (blue, red) compared to random simulations (black, gray), showing clear deviations from spatial randomness. **b**. Inflection point in the *α*1 distance curve between the fourth and the fifth neighbour generations, indicating structural organization into clusters. **c**. Schematic of the modeling algorithm incorporating observed point density, first nearest-neighbor distance, and empirical labeling efficiency to predict cluster sizes. **d**. Sum of squared errors heatmap for cluster models using uncorrected localization density. **e**. Sum of squared errors heatmap using corrected molecular density, demonstrating a lower error range and predicting a maximum cluster size between 13 and 21 subunits.

## Discussion

The spatial organization of membrane proteins regulates cellular function. Here, we used the high spatial resolution (localisation precision median 3-5 nm) of MINFLUX microscopy combined with DNA-PAINT technology, to generate quantitative maps of Na,K-ATPase organization in the plasma membrane of A498 cells. We directly visualized and identified Na,K-ATPase dimer configurations within the membrane, supporting the hypothesis of Na,K-ATPase self-oligomerization. The observed preferential nearest-neighbour distance of approximately 7 nm corresponds to the structural dimensions predicted for adjacent NKA complexes within a lipid bilayer, moving the field beyond statistical inference to direct structural validation.^12^

MINFLUX and DNA-PAINT successfully resolved individual Na,K-ATPase molecules within clusters, providing localization precision of a few nanometers. This precision distinguishes specific protein-protein interactions (like dimers) from random co-localization or labeling artifacts. The methodology implemented here, which includes the use of high-specificity nanobodies targeting EGFP-tagged Na,K-ATPase subunits and DNA-PAINT, ensures optimal labeling density and specificity crucial for quantitative spatial analysis.

These structural findings reinforce previous functional and biophysical evidence of Na,K-ATPase dimerization.^5,13^ Prior studies using FRET-FCS provided indirect evidence for dimers by detecting FRET-active complexes and calculating diffusion times. FRET-FCS measurements estimated that 34% ± 11% of the Na,K-ATPase population existed in a dimer configuration. Furthermore, the diffusion time for these FRET-active oligomers was 1.51 +/−0.18 times slower than the single-channel FCS curve (representing primarily monomers). This reduction aligns with the diffusion coefficient expected for dimers based on models of trans-membrane protein movement. Our MINFLUX-derived spatial maps provide structural confirmation that these previously characterized oligomers represent dimeric assemblies.^5^

To quantify the observed nanoscale organization, we refined traditional spatial statistics. Standard clustering analysis, such as Ripley’s K-function, confirms clustering relative to a random distribution but does not distinguish specific, closely associated dimers from unstructured aggregates. We implemented an analytical approach involving the sequential measurement of nearest-neighbour distances (first to n-th). Analyzing the distribution of the distance to the first nearest-neighbour and comparing it to higher-generation neighbours yields a quantitative metric of stoichiometry. A distinct increase in distance at the fifth and sixth neighbours provides statistical support for aggregation into complexes of up to 21 monomers (compensating for dark molecules), separating specific oligomerization from unspecific large-scale clustering, where the distances might increase more gradually. This approach enhances the capacity of nearest-neighbour analysis to decipher the stoichiometry of closely spaced protein complexes, enabled by the high localization precision in MINFLUX.

By examining the sequential neighbour distributions sequentially without restricting the regression to pre-defined template models, this methodology circumvents a constraint of the previously published template-matching SPINNA protocol, rendering it generalizable to complex target systems where native conformations are undetermined.^14^ To evaluate macromolecular stoichiometry without assuming a uniform cluster size based on raw coordinate inflections, the algorithm evaluates a continuous distribution of cluster sizes, modeled as a normal distribution between 2 and n subunits, while integrating observed molecular density. The mathematical model accounts for the undetected fraction of molecules resulting from incomplete labeling, incorporating the approximately 60% efficiency of anti-GFP nanobody binding under native conditions as recently characterized.^15^ Correcting for these unobserved dark states indicates that the biological assemblies have an average cluster size of 7 to 11 molecules. This framework serves as a complementary method to density-based segmentation algorithms such as DBSCAN. While DBSCAN isolates and classifies clusters based strictly on the visible point cloud, this approach incorporates experimental labeling efficiency to estimate the underlying biological distribution.

Quantifying these high-order architectures requires evaluating the interplay between transiently expressed exogenous fusion proteins and the resident pool of endogenous Na,K-ATPase complexes. Because the imaging workflow relies exclusively on anti-GFP nanobodies to target the EGFP-tagged constructs, the localizations capture only the exogenously introduced pool. In the native cellular environment, these labeled subunits must compete with endogenous *α* and *β* subunits for functional heterodimer assembly and biochemical trafficking to the cell surface. This competition introduces a molecular dilution factor. Consequently, the observed cluster sizes and stoichiometry obtained via spatial statistical modeling approach represent a conservative lower bound of the actual macromolecular arrangements. Stochastic incorporation of unlabeled endogenous subunits into these complexes means that the actual nanoclusters are likely larger and more complex than those explicitly resolved by MINFLUX localization alone. This competition indicates that the calculated presence of higher-order oligomers reflects a dense native environment.

The confirmed existence of Na,K-ATPase dimers and clusters suggests significant regulatory implications. Oligomerization potentially modulates enzymatic activity. A dimeric configuration may increase enzyme activity, by allowing two copies of the enzyme to operate asynchronously, mutually lowering the activation energy for ion transport, a mechanism demonstrated in the dimerization of sarco/endoplasmic reticulum Ca^2+^-ATPase (SERCA).^16^

Conversely, high-order clustering could lead to the opposite outcome. If Na,K-ATPase is concentrated within a small, diffusion-limited volume, competition for essential ligands such as ATP, Na^+^, and K^+^ could occur, reducing enzymatic activity. Therefore, the nanoscale organization, including dimerization and clustering, is likely a mechanism for both up- and down-regulation of enzymatic turnover. Recent super-resolution studies also indicate that *β* subunit glycosylation regulates this nanoscale organization by promoting stable clustering via galectin lattices.^7^

To validate the obtained localization maps, we evaluated potential artifacts arising from the labeling strategy. We could not resolve the two docking sites on the X2 nanobody, verifying that the observed dimerization is a biological feature rather than an artifact originating from the nanobody (*Supplementary Figure 4*). This technical confirmation provides a quality control step for the MINFLUX community utilizing similar labeling approaches.

We utilized 3D acquisitions despite the lower localization precision compared to 2D MINFLUX. Employing 2D imaging would superimpose the apical and basal membranes, potentially generating artificial clustering readouts and obscuring the native nanoscale organization of the protein (Supplementary Figure 5).

The membrane separation approach developed to address this 3D dimensionality revealed no differences between apical and basal clustering patterns for Na,K-ATPase, but it provides a computational tool for general imaging applications. This data-driven point cloud separation algorithm is applicable to 3D membrane datasets, including the analysis of adjacent membranes at cell contact sites, serving as an analytical method to mitigate clustering artifacts (*Supplementary Material*).

In conclusion, this study provides for the first time direct, high-resolution imaging evidence of Na,K-ATPase dimerization and nanoscale clustering in the plasma membrane, quantitatively confirmed via sequential nearest-neighbour analysis. Establishing 3D DNA-PAINT MIN-FLUX nanoscopy as a framework for structural verification enables future investigations, such as applying two-color MINFLUX tracking to monitor intramolecular conformational dynamics within single Na,K-ATPase molecules during enzymatic turnover, and visualizing the nanoscale proximity and modulation of Na,K-ATPase’s interaction with adaptor proteins and signaling partners.^17–20^

## Materials and Methods

### Cell Culture and Transfection

A498 cells (human kidney carcinoma) were grown at 37°C, 5% CO_2_ in RPMI 1640 Medium (Thermo Fisher Scientific) supplemented with 10% FBS (Thermo Fisher Scientific), 1% GlutaMAX (Thermo Fisher Scientific) and 1% Penicillin-Streptomycin solution (Thermo Fisher Scientific). Before seeding on the coverslip, cells were washed twice with PBS and incubated for 5 minutes with TripLE (Thermo Fisher Scientific) to facilitate detachment. After neutralization with cell culture medium, the suspension was centrifuged for 8 minutes at 3000 rpm, and the supernatant discarded. Cells were resuspended in the appropriate volume of complete cell culture media. Cells were seeded on 18 mm #1.5H coverslips (Marienfeld, # 0117580) in a 12-well plate (Corning) at 50% to 60% confluency, and incubated overnight. Upon reaching 70% to 80% confluency, cells were then transfected with Lipofectamine 3000 (Thermo Fisher Scientific) according to the manufacturer’s instructions. Briefly, 1 µg of either Na,K-ATPase*α*1-EGFP or Na,K-ATPase*β*1-EGFP (*Supplementary Material 2*) plasmid per well was diluted in Opti-MEM (Thermo Fisher Scientific), Lipofectamine, and P3000 mix according to the manufacturer’s instructions and distributed uniformly across the well. Transfection was carried out overnight before changing to fresh media, and then the cells were left to incubate for another 48 hours prior to fixation.

### DNA-PAINT Sample Preparation

Samples were fixed with pre-warmed 4% paraformaldehyde (PFA) (vol/vol in PBS, Electron Microscopy Sciences, #15710) for 10 minutes at room temperature and washed three times with PBS. Samples were then permeabilised with 0.5% (v/v) Triton X-100 in PBS for 5 minutes at room temperature, washed three times with PBS, and blocked with antibody incubation buffer (Massive Photonics) for 1 hour at room temperature under continuous agitation. Samples were then labelled with MASSIVE-TAG-X2 sdAB anti-GFP (two DNA docking sites, clone: 1H1 & 1B2, 5µM) nanobody at a ratio of 1:300 according to the manufacturer’s instructions, for one hour at room temperature. Samples were then washed three times with a 1X washing buffer (Massive Photonics) and incubated with 150 nm gold nanoparticles (Nanopartz # A11-150-CIT-DIH-1-25) for 3 minutes, followed by an extensive PBS wash. Coverslips were mounted on cavity slides (BMS # 12290) containing approximately 100 µl 0.5 nM solution of Abberior 660 MF3 imager strands diluted in imaging buffer (Massive Photonics) and sealed with dental glue (Picodent).

### 3D MINFLUX Imaging

Localisation events were acquired on an Abberior MINFLUX microscope (Abberior Instruments, Germany) using the 3D imaging sequence in the Imspector software (vol. m2205 and m2410). Single molecule events were acquired by 640 nm 3D donut excitation and targeted (split) fluorescence emission collected on two avalanche photodiodes, spectrally covering 650-680 nm and 685-720 nm. The 3D donut profile was optimized by adjusting a spatial light modulator (SLM) using 120 nm fluorescent beads (Abberior Instruments calibration sample). Biological calibration samples with nuclear pore complexes (NUP96-GFP U2OS cell line, Cytion 300174) were used to pre-calibrate excitation settings for DNA-PAINT MINFLUX (kit Abberior660). A high-numerical objective Olympus UPlanXApo 100x/1.40 Oil 8/0.17/FN26.5 was used for 3D localization.

To select imaging positions, transfected cells were identified using the EGFP signal by a 488 nm confocal scan (spectral detection 500-550 nm), adjusting the focal position close to the coverslip to capture the thin membrane sections of the cell. Small 3×3 µm MINFLUX ROIs were selected in a membrane area with stable gold bead coordinates (<0.2 nm in x,y; <1 nm in z). Prior to MINFLUX imaging, the binding dynamics of DNA-PAINT imagers were verified in an adjacent area using a 642 nm repeated confocal scan. MINFLUX Beamline Monitoring (MBM) was used for drift correction by picking six to seven in-focus 150 nm gold beads on the cover glass adjacent to the cell being imaged, which were revisited every 4 seconds. A pinhole size of 0.69 AU and 8-11% 642 nm laser power was used for all acquisitions. Each measurement was conducted for 2.5 to 3 hours per MINFLUX ROI and selected cell. Each ROI was selected from a different cell representing three biological replicates.

### Confocal Imaging

Confocal images were recorded using Zeiss LSM 980/Airyscan 2 point scanning system using a Plan-Apochromat 63x/1.4 oil DIC M27l objective. 16 bit z-stacks were recorded using bidirectional scanning with the pixel size and z-step optimized for Nyqvist sampling. A 488 nm laser was used to excite EGFP and the laser power was set to 1% with emission collected using a spectral detector in the 490 nm to 658 nm window. Gain was set to 760 V and the pinhole was set to 1 AU.

### Data Post Processing

Valid localisations were exported from Imspector Software as .npy files and loaded into PyMINFLUX for post-processing.^21^ Traces with fewer than 3 localisations were discarded, the reported localisation precision was derived by calculating the median σ for each dimension from localisations from traces with ≥ 3 localisations. Center frequency ratio (cfr) filtering was performed during the acquisition sequence and all localisations with cfr > 0.8 were considered invalid. When present, a secondary effective frequency (efo) peak was discarded, which usually corresponded to efo values exceeding 50 to 52 kHz. Refractive index mismatch was mitigated by multiplying all z coordinates by 0.7. To mitigate the reblinking artifact, trace id reassignment was performed by executing DBSCAN on all localization files (min pts = 1, eps = 3 nm), where detected clusters were assigned a trace id. After filtering, trace coordinates were calculated by averaging x,y, and z positions of remaining localisations and trace coordinates were exported as .csv files for further analysis.

### Membrane Fitting through Quadratic Programming

First, outliers were removed by density quantile thresholding and median absolute deviation (see Supplementary Material). The point pattern was then standardised. Initial point classification was determined by fitting a midplane (cubic polynomial) to the data via ridge regression, subject to regularisation parameter *α* = 0.001. Any points above the surface were classed as apical (and vice versa for basal). We denote the sets of data points labelled as apical and basal by the sets *A* and *B*, respectively. Then, we aim to minimise the mean square error (MSE) of two cubic polynomials (*f*_*a*_(*x, y*) and *f*_*b*_(*x, y*)) to the data, subject to a gap constraint (see Supplementary Material). Since *A* and *B* are disjoint sets, the surfaces are fitted to separate subsets of the data, which is analogous to minimising the following single-objective cost function

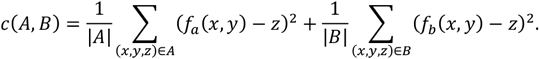

This reduces to a quadratic programming problem (QP), for which we may use one of three built-in solvers: Operator Splitting Quadratic Programming (OSQP), Embedded Conic Solver (ECOS) and Splitting Conic Solver (SCS). If all solvers fail, initial labels are used for classification. Once surfaces have been fit, points are re-classified by the surface they are closest to (i.e. apical or basal). In order to check for convergence, we calculate the total proportion of points which have changed class since the last iteration. If this value is below the tolerance parameter *tol*, we stop early, otherwise the process is repeated *max_iter* times. By default, we set *tol* = 0.01 and *max_iter* = 10. Once finished, outliers are returned to the point cloud and final classes are determined for each point based on which polynomial they lie closest to.

## Data Analysis

All data analysis was performed on averaged trace coordinates exported from PyMINFLUX. Briefly, point patterns were loaded into a Jupyter notebook and three non-overlapping 1×1 µm ROIs per dataset were isolated for further analysis. The median number of points per ROI was calculated and used for all simulations and all simulated point patterns had a minimum distance of 3 nm between points. k-NND curves were modelled by simulating 2D point patterns in 1×1µm space adjusting the cluster spatial size to the observed median first NND. The models probed clustering between 0-100% in 5% increments, and two different point densities, namely the observed median point density, and adjusted point density based on the known labeling efficiency. Data analysis scripts and MINFLUX imaging sequences can be found at github.com/bruno-stojcic/NKA-MINFLUX-Analysis. Data can be found at DOI: 10.71775/kth.vsztr-nq320

## Author Contributions

Hj.B. supervised the project. B.S. prepared the microscopy samples and performed the experiments. B.S., A.A, and H.B. performed MINFLUX acquisitions. B.S. implemented the data analysis. L.P. developed the membrane splitting algorithm. B.S. developed and implemented the cluster analysis algorithm. B.S. and A.A. produced the figures. B.S. and Hj.B. wrote the manuscript. All authors revised the manuscript.

## Acknowledgements

We acknowledge the ALM facility at SciLifeLab, KTH and the National Microscopy Infrastructure, NMI (VR-RFI 2023-00163) for providing assistance in microscopy.

L.P. acknowledges funding from the SciLifeLab RED Post-doctoral Fellowship (31005394).

Hj.B. acknowledges funding from the Swedish research council (VR) grant id: 2020-0534.

## Data Availability

The data is publicly available at DOI: 10.71775/kth.vsztr-nq320

## Notes

https://doi.org/10.71775/kth.vsztr-nq320

https://github.com/bruno-stojcic/NKA-MINFLUX-Analysis

### Competing Interest Statement

The authors have declared no competing interest.

