## Supplementary information for "Direct visualization of Na,K-ATPase clustering by 3D DNA-PAINT MINFLUX nanoscopy"

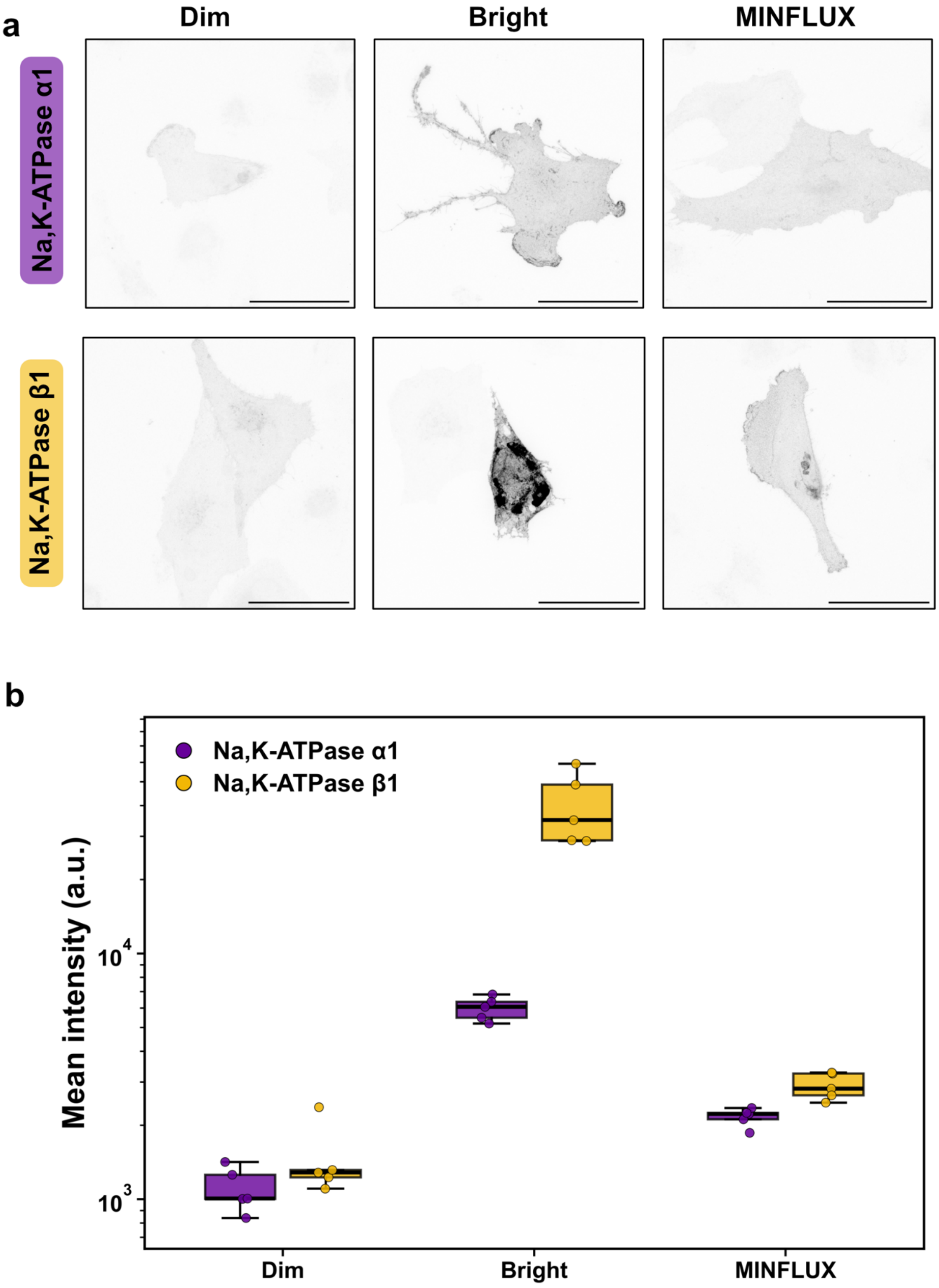
a
Dim
Bright
MINFLUX
Na,K-ATPase α1
Na,K-ATPase β1
b
Na,K-ATPase α1
Na,K-ATPase β1
Mean intensity (a.u.)
10^4
10^3
Dim
Bright
MINFLUX

**Supplementary Figure 1**: *Selection of cells for MINFLUX imaging based on transfection intensity.* **a.** Transfected samples exhibit heterogeneity in transfection efficiency, which is visible in the intensity and homogeneity of the GFP signal. Cells with low or high expression levels were excluded from MINFLUX imaging due to poor transfection efficiency or high intracellular endoplasmic reticulum signal accumulation. Labeled cells with excessive brightness also induced structural artifacts, including distorted cell morphology. Selected cells displayed a uniform GFP signal across the plasma membrane (scale bar: 50 μm). **b.** Circular regions of interest of 100x100 pixels were collected across maximum intensity projections of confocal z-stacks. Cell selection was quantified by measuring the background-subtracted mean gray value, and the log-scale plot defines the intensity range utilized to guide the selection of MINFLUX coordinates.

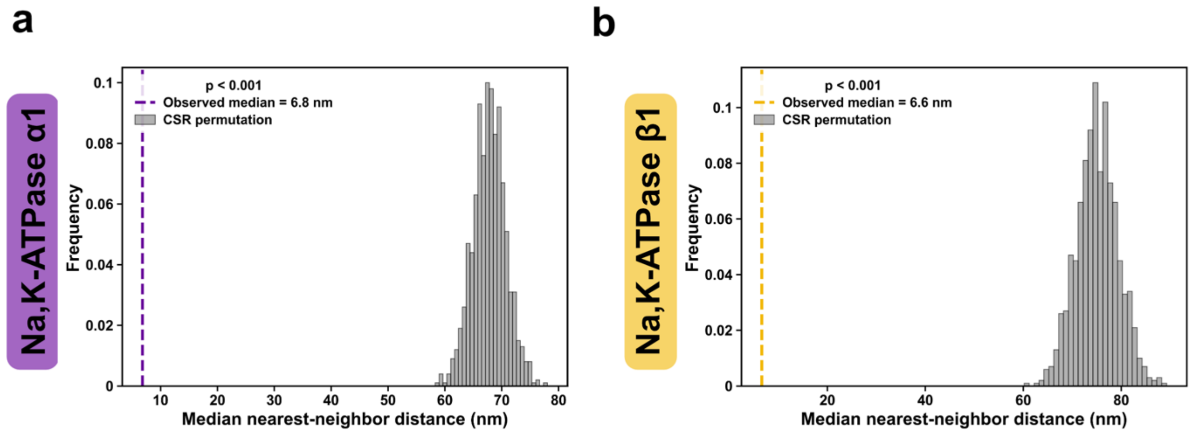


**Supplementary Figure 2**. *Permutation test displaying the distribution of medians for 3D complete spatial randomness simulations.* Complete spatial randomness (CSR) was simulated 1000 times within a 1 μm x 1 μm area using the observed axial distribution and the median number of observed coordinates per region of interest. The resulting permuted medians were compared to the experimental median using a Mann-Whitney U test. **a.** Distribution of simulated medians for the $\alpha 1$ subunit. **b.** Distribution of simulated medians for the $\beta 1$ subunit, where all comparisons indicate statistical significance ($p < 0.001$).

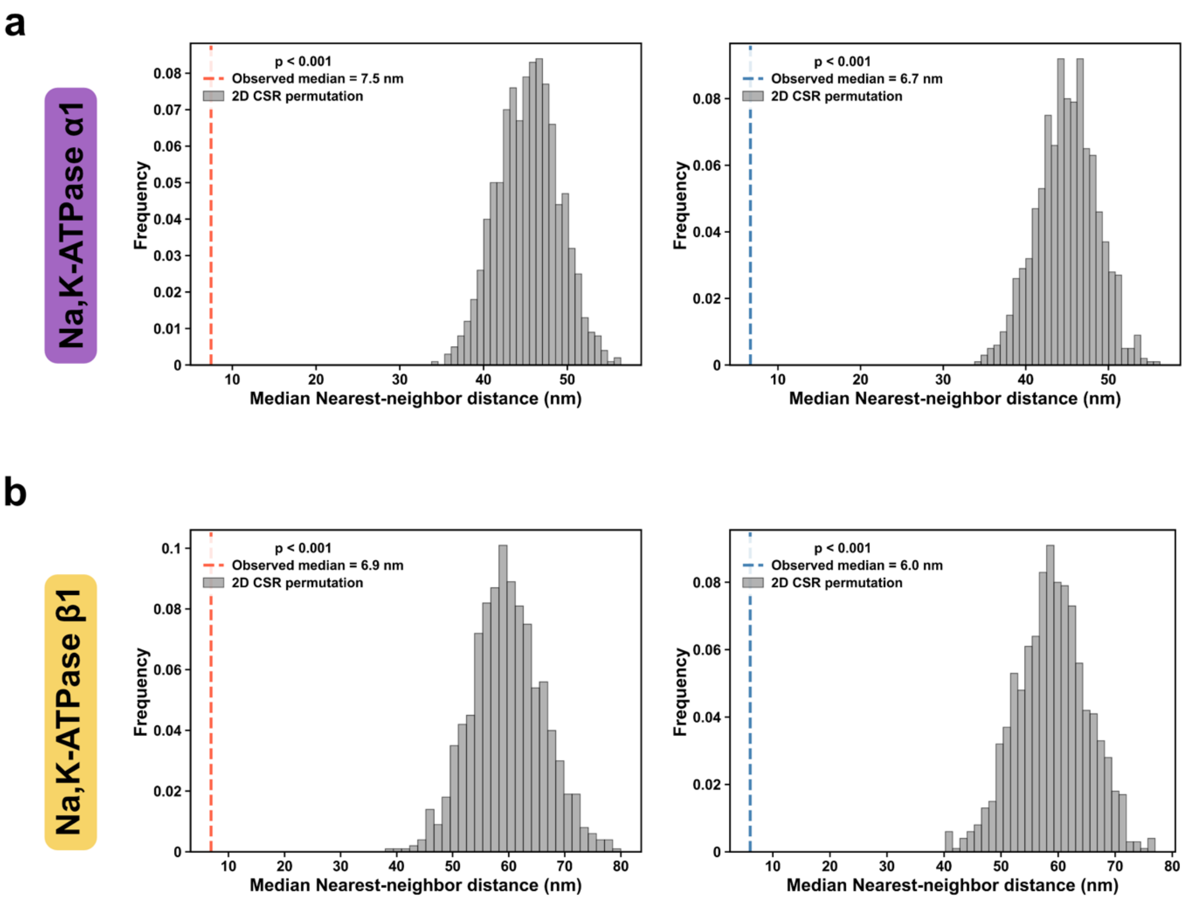


**Supplementary Figure 3**. *Permutation test displaying the distribution of medians for 2D complete spatial randomness.* Complete spatial randomness (CSR) was simulated 1000 times in a 1 μm x 1 μm coordinate space using the median number of detected points per region of interest, and permuted medians were compared against experimental values using a Mann-Whitney U test. **a.** Distribution of simulated medians for the $\alpha$1 subunit. **b.** Distribution of simulated medians for the $\beta$1 subunit, showing separate comparisons for the apical and basal membrane surfaces (p <0.001).

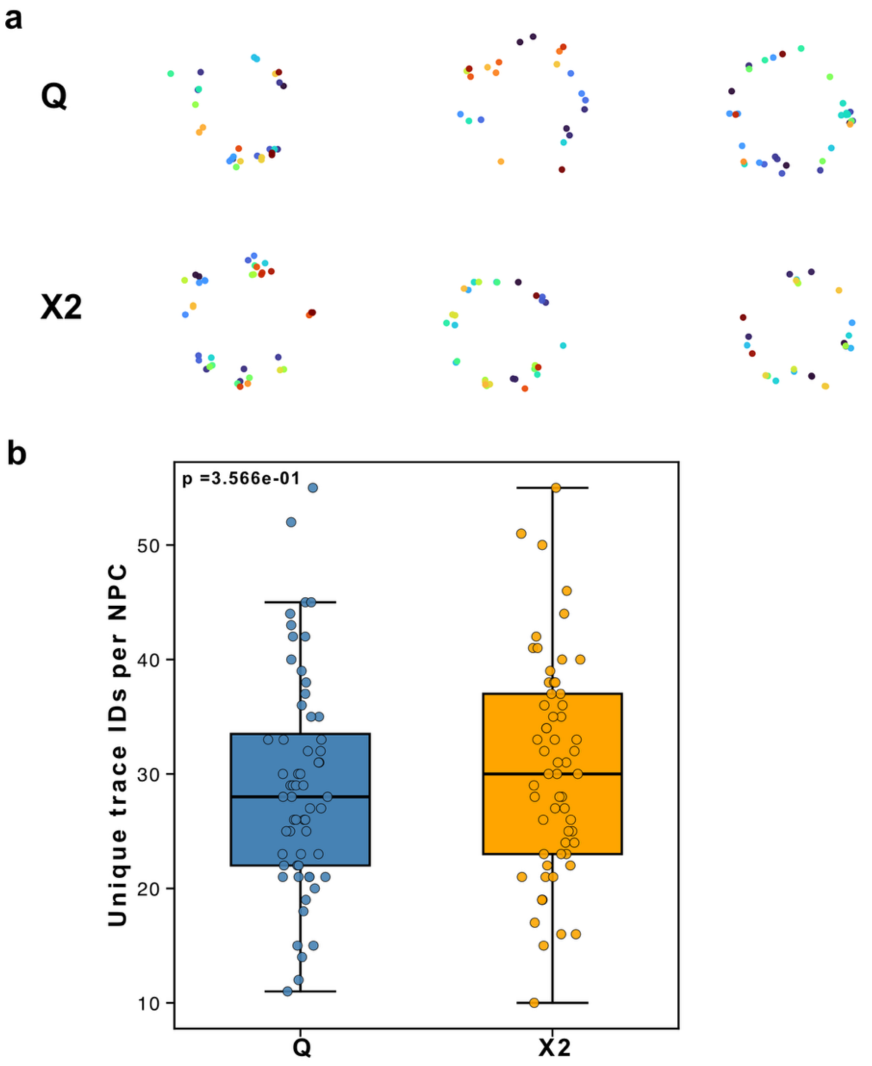


**Supplementary figure 5:** *Comparison of the 2D and 3D imaging of Nuclear Pore Complexes on MINFLUX*. **a.** Representative nuclear pore complexes recorded with 2D or 3D MINFLUX sequences, color-coded by trace ID. **b.** Quantification of unique trace IDs per nuclear pore complex over a 1-hour acquisition period across sixty complexes per condition. The 2D MINFLUX sequence yields a higher number of traces because targets from both sides of the nuclear envelope are projected onto a single lateral plane, necessitating axial resolution to distinguish true molecular positions. The increased trace count in 2D data is also attributed to the accelerated localization duty cycle of the 2D sequence compared to the 3D sequence. **c.** Lateral localization precision distribution for 2D acquisitions, demonstrating a precision approximately two-fold higher than the corresponding 3D sequence. **d** Localization precision distribution for 3D acquisitions. Although 3D imaging exhibits reduced lateral precision, the added axial resolution allows individual coordinates to be correctly assigned to their respective side of the nuclear envelope.

### Supplementary Material 1 – Membrane Fitting Algorithm

#### Outlier Removal

Outliers are removed via density quantile thresholding and median absolute deviation (MAD). For density quantile thresholding, the density of each point is estimated as the reciprocal of the mean distance to that point's closest 16 neighbours (by Euclidean distance) in the XY plane only. The points while lie within the bottom 2% of all densities are then classed as outliers. For MAD, the absolute difference in $z$ coordinate to the median $z$ coordinate is calculated for all points. Then, the median of all differences is determined. If any point has a difference from the median of more than 4 times the median of all differences, it is removed. Mathematically, all points which obey the following inequality are classed as outliers

$$\mathrm{med}(|z - \boldsymbol{z}|) \geq 4\,\mathrm{med}\big(\boldsymbol{z} - \mathrm{med}(|z - \boldsymbol{z}|)\big),$$

where med is the median function over vector space, $z$ is the point $z$ coordinate, and $\boldsymbol{z}$ is the vector of all $z$ coordinates. All points labelled as outliers are temporarily removed prior to fitting and classification (see below).

#### Gap Constraint

The gap constraint encourages the fitted polynomials to maintain some minimum distance along the z-axis. Either a hard constraint (forced) or soft constraint (encouraged) may be used. If not given, the gap size is set as $\Delta = 0.3(z_{0.95} - z_{0.05})$, where $z_{0.95}$ and $z_{0.05}$ are the $z$ coordinates in the 95$^{th}$ and 5$^{th}$ percentile of $\boldsymbol{z}$, respectively. The hard constraint imposes the inequality

$$f_a(x,y) - f_b(x,y) \geq \Delta\ \forall (x,y) \in A \cup B.$$

The soft constraint updates the cost function to

$$c(A,B;\Delta) = \frac{1}{|A|}\sum_{(x,y,z)\in A}(f_a(x,y) - z)^2 + \frac{1}{|B|}\sum_{(x,y,z)\in B}(f_b(x,y) - z)^2$$

$$+\lambda \max(\{\Delta - f_a(x,y) + f_b(x,y) \colon (x,y,z) \in A \cup B\} \cup \{0\})$$

where $\lambda$ is the user-defined weighting of the soft constraint. The hard constraint is used by default, with the soft constraint used as fallback in case of failure.

#### Data Simulation

Simulation parameters are described in **Table 1**, alongside the full range of possible parameter values. Each combination of parameter values was repeated 10 times, giving 781250 total simulations. First, a cubic polynomial representing the apical membrane, $h_a(x,y)$, is defined over the space $x \in [0, x_{max}]$, $y \in [0, y_{max}]$, with coefficients sampled randomly from a standard uniform distribution defined over [0, 1]. The coefficients of each term of the polynomial are rescaled such that their value at the edges of the domain could not exceed 1 (this prevents enormous weighting from higher order terms). For example, the coefficient of $x^2$ is divided by $x_{max}^2$ and the coefficient of $xy$ is divided by $x_{max}y_{max}$. We define $G$ as

$$G = \left\{\left(\frac{x_{max}n}{100}, \frac{y_{max}n}{100}\right): n \in \mathbb{N}, n \leq 100\right\},$$

representing a 100-by-100-point grid over the domain. The height of the polynomial is evaluated at all discrete points across $G$, and the minimum and maximum are taken as an approximation for global minimum and maximum, i.e. we assume

$$h_{min} = \min\big(h_a(x, y)\big) \approx \min(\{h(x, y): (x, y) \in G\}),$$

$$h_{max} = \max\big(h_a(x, y)\big) \approx \max(\{h(x, y): (x, y) \in G\}).$$

The polynomial undergoes an affine transformation to translate the approximate minimum to 0 and approximate maximum to $D_{max}$, that is

$$h_a(x, y) \coloneqq \frac{h_a(x, y) - h_{min}}{h_{max} - h_{min}} h_{max}.$$

Then, $N$ 2D data points are sampled randomly from a uniform distribution defined over the space $x' \in [0, 1]$, $y' \in [0, 1]$. These coordinates are augmented by the density augment function $g$ and rescaled back to $x \in [0, x_{max}]$, $y \in [0, y_{max}]$. The $z$ coordinate of each data point is calculated from the polynomial. This gives a set of $(x, y, z)$ coordinates, $C$, denoted baseline coordinates, where

$$C = \left\{\Big(x_{max}g(x), y_{max}g(y), h_a\big(x_{max}g(x), y_{max}g(y)\big)\Big): x, y \sim \mathrm{Unif}(0,1)\right\}.$$

For each baseline point, a 2D vector is sampled from a 2D Gaussian with mean at the baseline $(x, y)$ coordinate and standard deviation $\boldsymbol{\sigma}_{xy}$. A new $z$ coordinate is sampled from a 1D Gaussian distribution centred at the baseline $z$ coordinate with standard deviation $\sigma_z$. This gives the set of ground truth points associated with the apical membrane, $A_{true}$, where

$$A_{true} = \left\{(x_g, y_g, z_g): (x_g, y_g) \sim \mathrm{Norm}_2\left((x, y), \boldsymbol{\sigma}_{xy}\right), z_g \sim \mathrm{Norm}_1(z, \sigma_z), (x, y, z) \in C\right\},$$

with

$$\boldsymbol{\sigma}_{xy} = \begin{bmatrix} \sigma_{xy} & 0 \\ 0 & \sigma_{xy} \end{bmatrix}.$$

All points in $A_{true}$ are labelled as part of the apical membrane. For the basal membrane, a copy of the first polynomial is created and translated downward by the separation distance, $\Delta$, giving $h_b(x, y) = h_a(x, y) - \Delta$. The process of generating points is repeated as above, with new data points stored in the set $B_{true}$ and labelled as part of the basal membrane. Once all points have been generated, $\lceil 2Nq/100 \rceil$ additional outlier points are sampled randomly from a 3D uniform distribution defined between the minimum and maximum coordinates of all existing points. Outlier points are labelled as part of whichever polynomial they are closest to in Euclidean space.

| Parameter | Unit | Definition | Value(s) |
|---|---|---|---|
| $x_{max}$ | nm | Maximum size of $x$ domain. | 3000 |
| $y_{max}$ | nm | Maximum size of $y$ domain. | 3000 |
| $D_{max}$ | nm | Distance between minimum and maximum height of membrane. | (50, 100, 150, 200, 250) |
| $N$ | - | Number of points per membrane. | (500, 1000, 1500, 2000, 2500) |
| $g$ | - | Density augment function, used to update baseline coordinates and simulate density variation across the membrane. | $g(x) = x, x^2, x^3, x^4, x^5$ |
| $\sigma_{xy}$ | nm | The lateral spread of points from the membrane. | (5, 10, 15, 20, 25) |
| $\sigma_z$ | nm | The axial spread of points from the membrane. | (5, 10, 15, 20, 25) |
| $\Delta$ | nm | The expected gap, i.e. the minimum distance between the apical and basal membrane. | (20, 40, 60, 80, 100) |
| $q$ | % | The percentage of random outliers which are generated uniformly over the distribution. | (10, 20, 30, 40, 50) |

**Table 1:** Simulation parameters and the range of values considered.

#### Model Validation

Datasets are simulated as above. For each simulation, the ground truth classification (i.e. apical or basal membrane) is stored for each point. The separation algorithm is applied to all data sets to derive estimates of the apical and basal polynomials, $f_a$ and $f_b$, and to classify each point. Point classes are compared to the ground truth to assess model performance. The adjusted rand index (ARI) between the ground truth classes and labels found by the analysis herein is calculated as in the literature [1]. Root mean square error (RMSE), $R$, was calculated as,

$$R = \frac{1}{20000} \sum_{(x,y)\in G} \sqrt{(f_a(x,y) - h_a(x,y))^2} + \sqrt{(f_b(x,y) - h_b(x,y))^2}.$$

#### Hardware

Simulations were run in parallel across multiple CPU cores using PyTorch's built-in multiprocessing (see **Data Availability Statement** for code). Preliminary analysis (Figure A) was undertaken on a MacBook Pro Apple M3 with 8gb of RAM. Runtime was <5 seconds. Data simulation (Figure B) was performed on the Tetralith High Performance Computer (https://www.nsc.liu.se/systems/tetralith/) using 2x Intel Xeon Gold 6130 CPUs. 64 CPU cores were used with 192gb of RAM. All simulations were completed and analysed in ~40 minutes.

**Acknowledgements**

The computations and data handling were enabled by resources provided by the National Academic Infrastructure for Supercomputing in Sweden (NAISS), partially funded by the Swedish Research Council through grant agreement no. 2022-06725.

**Data Availability Statement**

The software package and data that support the findings of this study are available under the terms of the GNU General Public License v. 3.0 at: https://github.com/lucapanconi/separate_membranes.

**Data Availability**

The code and data which support the findings of this study are available at https://github.com/lucapanconi/separate_membranes.

**Justification**

**Outlier Removal**

The extreme height values of outliers can encourage the program to overfit. As such, outliers are first removed via density quantile thresholding and median absolute deviation (MAD). Since MAD relies on the median (as opposed to mean and standard deviation), it is not weighted by extreme values and is therefore robust to outliers.

**Ridge Regression**

Since we fit a cubic polynomial, the polynomial bases are inherently correlated and may express multi-collinearity. As such, we determine our initial fit through ridge regression, which uses a regularisation term to reduce the coefficients of larger bases, so that no one base dominates the others.

**Upper-Lower Membrane Simulation**

Since we aim to test the worst-case performance of the method, we assume all membranes lie a fixed vertical distance apart at all points. As such, we create a copy of the upper membrane translated downward by the separation distance, Δ.

| condition_name | density_label | total_points | cluster_size_n_max | cluster_fraction | cluster_percent | sigma_nm | sse |
|---|---|---|---|---|---|---|---|
| **Apical_2D** | detection_corrected_density | 188 | 19 | 0.6000000000000002 | 60.00000000000002 | 13.26171875 | 740.9125331993973 |
| **Apical_2D** | detection_corrected_density | 188 | 13 | 1.0000000000000002 | 100.00000000000003 | 13.274370193481445 | 792.3068399780773 |
| **Apical_2D** | detection_corrected_density | 188 | 13 | 0.9500000000000003 | 95.00000000000003 | 13.274370193481445 | 845.9710844196528 |
| **Apical_2D** | detection_corrected_density | 188 | 20 | 0.6000000000000002 | 60.00000000000002 | 14.803306579589844 | 846.7298587174967 |
| **Apical_2D** | detection_corrected_density | 188 | 18 | 0.6500000000000001 | 65.00000000000001 | 13.26171875 | 863.4610661361608 |
| **Apical_2D** | detection_corrected_density | 188 | 14 | 0.9000000000000002 | 90.00000000000003 | 13.274370193481445 | 885.1950185759099 |
| **Apical_2D** | detection_corrected_density | 188 | 15 | 0.8000000000000003 | 80.00000000000003 | 13.455078125 | 892.514388795232 |
| **Apical_2D** | detection_corrected_density | 188 | 14 | 0.9500000000000003 | 95.00000000000003 | 13.274370193481445 | 914.9752683826042 |
| **Apical_2D** | detection_corrected_density | 188 | 17 | 0.7000000000000002 | 70.00000000000001 | 13.088327884674072 | 925.6770889120105 |
| **Apical_2D** | detection_corrected_density | 188 | 21 | 0.5500000000000002 | 55.000000000000014 | 14.803306579589844 | 928.0099199615022 |

**Supplementary Table 2:** Top 10 best fitting k-NND models for Na,K-ATPase $\alpha$1 apical membrane.

| condition_name | density_label | total_points | cluster_size_n_max | cluster_fraction | cluster_percent | sigma_nm | sse |
|---|---|---|---|---|---|---|---|
| **Basal_2D** | detection_corrected_density | 192 | 20 | 0.5500000000000002 | 55.000000000000014 | 6.6875 | 56.0278044195403 |
| **Basal_2D** | detection_corrected_density | 192 | 21 | 0.5500000000000002 | 55.000000000000014 | 6.6875 | 136.46929788033574 |
| **Basal_2D** | detection_corrected_density | 192 | 13 | 0.9500000000000003 | 95.00000000000003 | 10.85076904296875 | 175.54037238096328 |
| **Basal_2D** | detection_corrected_density | 192 | 13 | 1.0000000000000002 | 100.00000000000003 | 10.85076904296875 | 184.639209258829 |
| **Basal_2D** | detection_corrected_density | 192 | 14 | 0.8500000000000003 | 85.00000000000003 | 10.85076904296875 | 206.68779868574373 |
| **Basal_2D** | detection_corrected_density | 192 | 13 | 0.9000000000000002 | 90.00000000000003 | 10.85076904296875 | 219.0817272302087 |
| **Basal_2D** | detection_corrected_density | 192 | 14 | 0.9000000000000002 | 90.00000000000003 | 10.85076904296875 | 299.1338666243291 |
| **Basal_2D** | detection_corrected_density | 192 | 12 | 1.0000000000000002 | 100.00000000000003 | 9.587890625 | 299.95295966602055 |
| **Basal_2D** | detection_corrected_density | 192 | 14 | 0.8000000000000003 | 80.00000000000003 | 10.85076904296875 | 302.164739181134 |
| **Basal_2D** | detection_corrected_density | 192 | 17 | 0.6500000000000001 | 65.00000000000001 | 6.6875 | 344.30496382524444 |

**Supplementary Table 3:** Top 10 best fitting k-NND models for Na,K-ATPase $\alpha$1 basal membrane.

### Supplementary Material 2 – Plasmid Sequences

#### Na,K-ATPase B1-EGFP

TAGAATGCAGTGAAAAAAATGCTTTATTTGTGAAATTTGTGATGCTATTGCTTTATTTGTAAC
CATTATAAGCTGCAATAAACAAGTTAACAACAACAATTGCATTCATTTTATGTTTCAGGTTCA
GGGGGAGGTGTGGGAGGTTTTTTAAAGCAAGTAAAACCTCTACAAATGTGGTATGGCTGAT
TATGATCAGTTATCTAGAACTAGTGGATCCCCCGGGCTGCAGGAATTCGATCCATAGAGCT
ACATGCTTCATTCCAGGACGTCTTGCTTCCCCACATGCTGCGGTGCTTTCCTACCAGGGTA
GAGTTCCAAACTCCAAGACTGAAGTACACAAAGAGGGGGTGGGTGTCGGATGCAGAGTGT
GTGGCCTGATGCTCCACGGCGTGCAGGACGGGGGGCTAATAGTAGGTTTCCTTCTCCACC
CAGCCGCCAGGGCGTCGCCTGATGATGAGTTTTCTGACTTCGTCATATACGAAGATGAGAA
GAGAGTAGGGGAAGGCACAGAACCACCAGGTAGGTTTGAGGGGATACATCCTAAGAGCAA
CACCCATTCCAGGGCAGTAGGAAAGGAAAGCAGCCAGGGCTGTCTCTTCAAAGAGGCCAA
ATATCAAGATCTTGTTCTTCATCCCCTGCTGGAAGACCGAATTCCTCCTGGTCTTACAGATG
ACCAAGTCGGCCCACTGCACCACCACGATACTGACGAAGAAGGCTGTGTGGCAGGTGAAC
TCCACGATTTTCCTCTGCTCATAGGTCCACTGCTGCCCGTAGCTGTCTTCCACATCGTTGA
TCCAGCGGTCATCCCAGTCCACTCGGAGGCCCAACAGGTGAATTGGGAGGAAGCCGTTCT
CAGCCAGAATCACAAAGTAAGTAAAGAAGCCTCCCAGGGCCTGGATCATTCCAATCTGCCC
ATAGGCCATGCTGATCAGCCGCTCATTCACAAGTTTGTCTGTTTTGGGATTTCTGGGCTGT
CTCTTCATGATGTCACTCTCAGCCTGCTCATAAGCCAGGGAGATGGCAGGAACCATGTCAG
TGCCCAAGTCAATGCAGAGGATGGTGACAGTCCCCAGTGGTAGTGGAATGTTTGCAATAAT
AAATATCAGGAACGGGGTGATCTCGGGAATGTTACTGGTTAAGGTATAAGCAATGGATTTC
TTCAAGTTATCAAAGATCAGACGACCTTCCTCTACTCCAGTCACAATTGAGGCAAAGTTGTC
ATCCAGAAGAATCATGTCAGCAGCTTGCTTGGACACATCTGAGCCAGCAATCCCCATAGCA
ACCCCAATGTCTGCTTTCTTCAAAGCTGGAGAGTCATTCACACCGTCACCAGTCACAGCCA
CGATAGCACCCTGTCTTTGGCAGCCTTCCACAATGATGAGCTTCTGCTGAGGGGAGGTCC
TGGCAAACACTATCTCAGTGTGGTACTTCAAAATGTCATCCAGCTGCTCGGAGGTCATGTC
CTTTAGATCACTGCCGTGTACTACGCAGGCCTTGGCATCCCTGGGGTTCACCTGGCTGAC
TGGGATGTTGAGGCGGGCAGCAATGTCTTCCACGGTCTCATTGCCTTCTGAGATGATGCC
CACACCTTTGGCAATAGCTTTAGCTGTGATTGGATGGTCTCCTGTGACCATGATGACCTTA
ATTCCAGCACTTCGACATTTGCCCACGGCATCAGGAACGGCCGCCCGTGGAGGGTCAATC
ATGGAGATGAGCCCAACAAAGCACAGATTATCGATAGGGAAATTCACATCGTCAGTGTCAA
ACTGGAACCCTTCAGGAAACTGTTCATCTGGCAGAAAGAGGTGGCAGAAACCTAGGACTC
GTTCTCCGAGGCCCCCCAGCTCCAAATAGGCGTTCTGAAAGGCGTCTTTCAGCTCCTCATC
CAGGGGCTGCTCCTTGCCGTGGAGGAGGATAGAGCTGCAACGGTCTAGGATCCTTTCTGG
GGCGCCCTTCATCACCAACAGGTGTTGGGGCTCCGATGTGTTGGGGTTCTTATGAATAGA
CAACTGGTACTTGTTGGTGGAGTTGAAGGGTATCTCGACGATTTTGGCGTATCTTTCTCTC
ATCTCCTTCACGGAACCACAGCACAGCTCTATGCACTTTAAGAGTGCTGACTCAGAGGCAT
CTCCTGCAACTGCCCGCTTAAGAATAGGTAGGTTTTCCTGGTTAGCCTGAAACACTGCCCT
GTTACAAAGACCTGCAATTCTGGACAGAGCAAGCCAGGTAGCTGAAGTCTTGTCAAAAGAG
ACACCACTCTGATTCTCTGTCGTATCAGCTTCATGGATTTGATTGTCAAACCACATGTGGGC
CACTGTCATCCGGTTCTGAGTCAGAGTTCCAGTTTTATCAGAGCAGATGGTGGACGTGGAC
CCCAAGGTCTCCACAGCTTCTAAGTTCTTCACTAAGCAGTTTTTCCTTGCCATGCGTTTGGC
AGTAAGTGTCAGACAGACCGTGACAGTGGCCAGCAAACCTTCCGGCACATTGGCTACGAT
GATACCGATGAGGAAGATGACAGCCTCAAGCCAGGTGTACTCAAGGATGAGAGAAAGGAT
GAAGAAAGACACACCCAGGAACACAGCCACACCCGTGATGATGTGGATAAAATGTTCAATT
TCTGCAGCAATGGGGGTCTGGCCTCCTTCCAGCCCAGAAGCAAGTGTGGCAATTTTTCCC
ATCACAGTGCGATCCCCAGTGTAGACAACAATACCACGTGCGGTGCCTTCAACACAATTGG
TTGAAAAGAAGGCAATGTTCCTCGTCTCCAGGGGGTTTTCATTTGTGAAATCTGGAGACCT
AGTCTGGGGTTCTGATTCACCGGTGAGCGAGGAGTTATCCACCTTGCAGCCATTTGCAGAT
ATGATTCTGAGGTCAGCAGGAATTCGGTCTCCTCCTTTTACTTCCACCAGATCCCCAACCA

CAACTTCCTCCGCATTTATGCTCATTTTCTCACCATTTCGAATCACAAGGGCTTGCTGAGGG
ACCATGTTTTTGAAGGATTCCATGATCTTTGAACTTTTAGCTTCTTGATAGTAGGAGAAGCA
ACCAGTTATGATTACAACGGCTGATAGCACCACACCCAGGTACAGATTATCGTTTTGAGGT
TCCTCTTCTGTAGCAGCTTGGATGCTATAAGCCAAGAAACAAAGAATCGCTCCAATCCACA
GTAACATTGAGAACCCCCCAAAGAGCTGCCGACAAAACTTGATCCATTCAGGAGTAGTGG
GAGGGGGAGTGAGGGCGTTGGGACCATCTCGCGCCAGGATCTCAGCTGCACGAGCAGAT
GTTAATCCCCGGCTCAAGTCTGTTCCATATTTACGATGAAGTTCATCAAGGCTAAGTTTATG
ATCATCCATAGAAACTTCTTTCTTCAGTTCATCCATGTCCCTGTCTTTTTTGCCCTTTTTGCC
CTTTTTATCACCTTGTTCTGAAACAGCTGCAGGCTCATACTTATCACGTCCCGAAGCTTGAG
CTCGAGATCTGAGTCCGGACTTGTACAGCTCGTCCATGCCGAGAGTGATCCCGGCGGCG
GTCACGAACTCCAGCAGGACCATGTGATCGCGCTTCTCGTTGGGGTCTTTGCTCAGGGCG
GACTGGGTGCTCAGGTAGTGGTTGTCGGGCAGCAGCACGGGGCCGTCGCCGATGGGGG
TGTTCTGCTGGTAGTGGTCGGCGAGCTGCACGCTGCCGTCCTCGATGTTGTGGCGGATCT
TGAAGTTCACCTTGATGCCGTTCTTCTGCTTGTCGGCCATGATATAGACGTTGTGGCTGTT
GTAGTTGTACTCCAGCTTGTGCCCCAGGATGTTGCCGTCCTCCTTGAAGTCGATGCCCTTC
AGCTCGATGCGGTTCACCAGGGTGTCGCCCTCGAACTTCACCTCGGCGCGGGTCTTGTAG
TTGCCGTCGTCCTTGAAGAAGATGGTGCGCTCCTGGACGTAGCCTTCGGGCATGGCGGAC
TTGAAGAAGTCGTGCTGCTTCATGTGGTCGGGGTAGCGGCTGAAGCACTGCACGCCGTAG
GTCAGGGTGGTCACGAGGGTGGGCCAGGGCACGGGCAGCTTGCCGGTGGTGCAGATGA
ACTTCAGGGTCAGCTTGCCGTAGGTGGCATCGCCCTCGCCCTCGCCGGACACGCTGAACT
TGTGGCCGTTTACGTCGCCGTCCAGCTCGACCAGGATGGGCACCACCCCGGTGAACAGC
TCCTCGCCCTTGCTCACCATGGTGGCGACCGGTAGCGCTAGCGGATCTGACGGTTCACTA
AACCAGCTCTGCTTATATAGACCTCCCACCGTACACGCCTACCGCCCATTTGCGTCAATGG
GGCGGAGTTGTTACGACATTTTGGAAAGTCCCGTTGATTTTGGTGCCAAAACAAACTCCCA
TTGACGTCAATGGGGTGGAGACTTGGAAATCCCCGTGAGTCAAACCGCTATCCACGCCCA
TTGATGTACTGCCAAAACCGCATCACCATGGTAATAGCGATGACTAATACGTAGATGTACT
GCCAAGTAGGAAAGTCCCATAAGGTCATGTACTGGGCATAATGCCAGGCGGGCCATTTAC
CGTAAGTTATGTAACGCGGAACTCCATATATGGGCTATGAACTAATGACCCCGTAATTGATT
ACTATTAATAACTAATGCATGGCGGTAATACGGTTATCCACAGAATCAGGGGATAACGCAG
GAAAGAACATGTGAGCAAAAGGCCAGCAAAAGGCCAGGAACCGTAAAAAGGCCGCGTTGC
TGGCGTTTTTCCATAGGCTCCGCCCCCCTGACGAGCATCACAAAAATCGACGCTCAAGTCA
GAGGTGGCGAAACCCGACAGGACTATAAAGATACCAGGCGTTTCCCCCTGGAAGCTCCCT
CGTGCGCTCTCCTGTTCCGACCCTGCCGCTTACCGGATACCTGTCCGCCTTTCTCCCTTCG
GGAAGCGTGGCGCTTTCTCATAGCTCACGCTGTAGGTATCTCAGTTCGGTGTAGGTCGTTC
GCTCCAAGCTGGGCTGTGTGCACGAACCCCCCGTTCAGCCCGACCGCTGCGCCTTATCC
GGTAACTATCGTCTTGAGTCCAACCCGGTAAGACACGACTTATCGCCACTGGCAGCAGCC
ACTGGTAACAGGATTAGCAGAGCGAGGTATGTAGGCGGTGCTACAGAGTTCTTGAAGTGG
TGGCCTAACTACGGCTACACTAGAAGAACAGTATTTGGTATCTGCGCTCTGCTGAAGCCAG
TTACCTTCGGAAAAAGAGTTGGTAGCTCTTGATCCGGCAAACAAACCACCGCTGGTAGCG
GTGGTTTTTTTGTTTGCAAGCAGCAGATTACGCGCAGAAAAAAAGGATCTCAAGAAGATCC
TTTGATCTTTTCTACGGGGTCTGACGCTCAGTGGAACGAAAACTCACGTTAAGGGATTTTG
GTCATGAGATTATCAAAAAGGATCTTCACCTAGATCCTTTTAAATTAAAAATGAAGTTTTAAA
TCAATCTAAAGTATATATGAGTAACCTGAGGCTATGGCAGGGCCTGCCGCCCCGACGTTG
GCTGCGAGCCCTGGGCCTTCACCCGAACTTGGGGGGTGGGGTGGGGAAAAGGAAGAAAC
GCGGGCGTATTGGCCCCAATGGGGTCTCGGTGGGGTATCGACAGAGTGCCAGCCCTGGG
ACCGAACCCCGCGTTTATGAACAAACGACCCAACACCGTGCGTTTTATTCTGTCTTTTTATT
GCCGTCATAGCGCGGGTTCCTTCCGGTATTGTCTCCTTCCGTGTTTCAGTTAGCCTCCCCC
TAGGGTGGGCGAAGAACTCCAGCATGAGATCCCCGCGCTGGAGGATCATCCAGCCGGCG
TCCCGGAAAACGATTCCGAAGCCCAACCTTTCATAGAAGGCGGCGGTGGAATCGAAATCT
CGTGATGGCAGGTTGGGCGTCGCTTGGTCGGTCATTTCGAACCCCAGAGTCCCGCTCAGA
AGAACTCGTCAAGAAGGCGATAGAAGGCGATGCGCTGCGAATCGGGAGCGGCGATACCG

TAAAGCACGAGGAAGCGGTCAGCCCATTCGCCGCCAAGCTCTTCAGCAATATCACGGGTA
GCCAACGCTATGTCCTGATAGCGGTCCGCCACACCCAGCCGGCCACAGTCGATGAATCCA
GAAAAGCGGCCATTTTCCACCATGATATTCGGCAAGCAGGCATCGCCATGGGTCACGACG
AGATCCTCGCCGTCGGGCATGCTCGCCTTGAGCCTGGCGAACAGTTCGGCTGGCGCGAG
CCCCTGATGCTCTTCGTCCAGATCATCCTGATCGACAAGACCGGCTTCCATCCGAGTACGT
GCTCGCTCGATGCGATGTTTCGCTTGGTGGTCGAATGGGCAGGTAGCCGGATCAAGCGTA
TGCAGCCGCCGCATTGCATCAGCCATGATGGATACTTTCTCGGCAGGAGCAAGGTGAGAT
GACAGGAGATCCTGCCCCGGCACTTCGCCCAATAGCAGCCAGTCCCTTCCCGCTTCAGTG
ACAACGTCGAGCACAGCTGCGCAAGGAACGCCCGTCGTGGCCAGCCACGATAGCCGCGC
TGCCTCGTCTTGCAGTTCATTCAGGGCACCGGACAGGTCGGTCTTGACAAAAAGAACCGG
GCGCCCCTGCGCTGACAGCCGGAACACGGCGGCATCAGAGCAGCCGATTGTCTGTTGTG
CCCAGTCATAGCCGAATAGCCTCTCCACCCAAGCGGCCGGAGAACCTGCGTGCAATCCAT
CTTGTTCAATCATGCGAAACGATCCTCATCCTGTCTCTTGATCGATCTTTGCAAAAGCCTAG
GCCTCCAAAAAAGCCTCCTCACTACTTCTGGAATAGCTCAGAGGCCGAGGCGGCCTCGGC
CTCTGCATAAATAAAAAAAATTAGTCAGCCATGGGGCGGAGAATGGGCGGAACTGGGCGG
AGTTAGGGGCGGGATGGGCGGAGTTAGGGGCGGGACTATGGTTGCTGACTAATTGAGAT
GCATGCTTTGCATACTTCTGCCTGCTGGGGAGCCTGGGGACTTTCCACACCTGGTTGCTG
ACTAATTGAGATGCATGCTTTGCATACTTCTGCCTGCTGGGGAGCCTGGGGACTTTCCACA
CCCTAACTGACACACATTCCACAGCTGGTTCTTTCCGCCTCAGGACTCTTCCTTTTTCAATA
TTATTGAAGCATTTATCAGGGTTATTGTCTCATGAGCGGATACATATTTGAATGTATTTAGAA
AAATAAACAAATAGGGGTTCCGCGCACATTTCCCCGAAAAGTGCCACCTGACGCGCCCTG
TAGCGGCGCATTAAGCGCGGCGGGTGTGGTGGTTACGCGCAGCGTGACCGCTACACTTG
CCAGCGCCCTAGCGCCCGCTCCTTTCGCTTTCTTCCCTTCCTTTCTCGCCACGTTCGCCGG
CTTTCCCCGTCAAGCTCTAAATCGGGGGCTCCCTTTAGGGTTCCGATTTAGTGCTTTACGG
CACCTCGACCCCAAAAAACTTGATTAGGGTGATGGTTCACGTAGTGGGCCATCGCCCTGAT
AGACGGTTTTTCGCCCTTTGACGTTGGAGTCCACGTTCTTTAATAGTGGACTCTTGTTCCAA
ACTGGAACAACACTCAACCCTATCTCGGTCTATTCTTTTGATTTATAAGGGATTTTGCCGAT
TTCGGCCTATTGGTTAAAAAATGAGCTGATTTAACAAAAATTTAACGCGAATTTTAACAAAAT
ATTAACGCTTACAATTTACGCGTTAAGATACATTGATGAGTTTGGACAAACCACAAC

**Na,K-ATPase B1-EGFP**
GTTTTCGTTCCACTGAGCGTCAGACCCCGTAGAAAAGATCAAAGGATCTTCTTGAGATCCT
TTTTTTCTGCGCGTAATCTGCTGCTTGCAAACAAAAAAACCACCGCTACCAGCGGTGGTTT
GTTTGCCGGATCAAGAGCTACCAACTCTTTTTCCGAAGGTAACTGGCTTCAGCAGAGCGCA
GATACCAAATACTGTTCTTCTAGTGTAGCCGTAGTTAGGCCACCACTTCAAGAACTCTGTAG
CACCGCCTACATACCTCGCTCTGCTAATCCTGTTACCAGTGGCTGCTGCCAGTGGCGATAA
GTCGTGTCTTACCGGGTTGGACTCAAGACGATAGTTACCGGATAAGGCGCAGCGGTCGGG
CTGAACGGGGGGTTCGTGCACACAGCCCAGCTTGGAGCGAACGACCTACACCGAACTGA
GATACCTACAGCGTGAGCTATGAGAAAGCGCCACGCTTCCCGAAGGGAGAAAGGCGGAC
AGGTATCCGGTAAGCGGCAGGGTCGGAACAGGAGAGCGCACGAGGGAGCTTCCAGGGG
GAAACGCCTGGTATCTTTATAGTCCTGTCGGGTTTCGCCACCTCTGACTTGAGCGTCGATT
TTTGTGATGCTCGTCAGGGGGGCGGAGCCTATGGAAAAACGCCAGCAACGCGGCCTTTTT
ACGGTTCCTGGCCTTTTGCTGGCCTTTTGCTCACATGTTCTTTCCTGCGTTATCCCCTGATT
CTGTGGATAACCGTATTACCGCCATGCATTAGTTATTAATAGTAATCAATTACGGGGTCATT
AGTTCATAGCCCATATATGGAGTTCCGCGTTACATAACTTACGGTAAATGGCCCGCCTGGC
TGACCGCCCAACGACCCCCGCCCATTGACGTCAATAATGACGTATGTTCCCATAGTAACGC
CAATAGGGACTTTCCATTGACGTCAATGGGTGGAGTATTTACGGTAAACTGCCCACTTGGC
AGTACATCAAGTGTATCATATGCCAAGTACGCCCCCTATTGACGTCAATGACGGTAAATGG
CCCGCCTGGCATTATGCCCAGTACATGACCTTATGGGACTTTCCTACTTGGCAGTACATCT
ACGTATTAGTCATCGCTATTACCATGGTGATGCGGTTTTGGCAGTACATCAATGGGCGTGG
ATAGCGGTTTGACTCACGGGGATTTCCAAGTCTCCACCCCATTGACGTCAATGGGAGTTTG
TTTTGGCACCAAAATCAACGGGACTTTCCAAAATGTCGTAACAACTCCGCCCCATTGACGC
AAATGGGCGGTAGGCGTGTACGGTGGGAGGTCTATATAAGCAGAGCTGGTTTAGTGAACC
GTCAGATCCGCTAGCGCTACCGGTCGCCACCATGGTGAGCAAGGGCGAGGAGCTGTTCA
CCGGGGTGGTGCCCATCCTGGTCGAGCTGGACGGCGACGTAAACGGCCACAAGTTCAGC
GTGTCCGGCGAGGGCGAGGGCGATGCCACCTACGGCAAGCTGACCCTGAAGTTCATCTG
CACCACCGGCAAGCTGCCCGTGCCCTGGCCCACCCTCGTGACCACCCTGACCTACGGCG
TGCAGTGCTTCAGCCGCTACCCCGACCACATGAAGCAGCACGACTTCTTCAAGTCCGCCA
TGCCCGAAGGCTACGTCCAGGAGCGCACCATCTTCTTCAAGGACGACGGCAACTACAAGA
CCCGCGCCGAGGTGAAGTTCGAGGGCGACACCCTGGTGAACCGCATCGAGCTGAAGGGC
ATCGACTTCAAGGAGGACGGCAACATCCTGGGGCACAAGCTGGAGTACAACTACAACAGC
CACAACGTCTATATCATGGCCGACAAGCAGAAGAACGGCATCAAGGTGAACTTCAAGATCC
GCCACAACATCGAGGACGGCAGCGTGCAGCTCGCCGACCACTACCAGCAGAACACCCCC
ATCGGCGACGGCCCCGTGCTGCTGCCCGACAACCACTACCTGAGCACCCAGTCCGCCCT
GAGCAAAGACCCCAACGAGAAGCGCGATCACATGGTCCTGCTGGAGTTCGTGACCGCCG
CCGGGATCACTCTCGGCATGGACGAGCTGTACAAGTCCGGACTCAGATCTCGAGCTCAAG
CTTCGAATTCCATGGCCCGCGGGAAAGCCAAGGAGGAGGGCAGCTGGAAGAAATTCATCT
GGAACTCAGAGAAGAAGGAGTTTCTGGGCAGGACCGGTGGCAGTTGGTTTAAGATCCTTC
TATTCTACGTAATATTTTATGGCTGCCTGGCTGGCATCTTCATCGGAACCATCCAAGTGATG
CTGCTCACCATCAGTGAATTTAAGCCCACATATCAGGACCGAGTGGCCCCGCCAGGATTAA
CACAGATTCCTCAGATCCAGAAGACTGAAATTTCCTTTCGTCCTAATGATCCCAAGAGCTAT
GAGGCATATGTACTGAACATAGTTAGGTTCCTGGAAAAGTACAAAGATTCAGCCCAGAGGG
ATGACATGATTTTTGAAGATTGTGGCGATGTGCCCAGTGAACCGAAAGAACGAGGAGACTT
TAATCATGAACGAGGAGAGCGAAAGGTCTGCAGATTCAAGCTTGAATGGCTGGGAAATTG

CTCTGGATTAAATGATGAAACTTATGGCTACAAAGAGGGCAAACCGTGCATTATTATAAAGC
TCAACCGAGTTCTAGGCTTCAAACCTAAGCCTCCCAAGAATGAGTCCTTGGAGACTTACCC
AGTGATGAAGTATAACCCAAATGTCCTTCCCGTTCAGTGCACTGGCAAGCGAGATGAAGAT
AAGGATAAAGTTGGAAATGTGGAGTATTTTGGACTGGGCAACTCCCCTGGTTTTCCTCTGC
AGTATTATCCGTACTATGGCAAACTCCTGCAGCCCAAATACCTGCAGCCCCTGCTGGCCGT
ACAGTTCACCAATCTTACCATGGACACTGAAATTCGCATAGAGTGTAAGGCGTACGGTGAG
AACATTGGGTACAGTGAGAAAGACCGTTTTCAGGGACGTTTTGATGTAAAAATTGAAGTTAA
GAGCTGATCACAAGCACAAATCTTTCCCACGGTACCGCGGGCCCGGGATCCACCGGATCT
AGATAACTGATCATAATCAGCCATACCACATTTGTAGAGGTTTTACTTGCTTTAAAAAACCTC
CCACACCTCCCCCTGAACCTGAAACATAAAATGAATGCAATTGTTGTTGTTAACTTGTTTAT
TGCAGCTTATAATGGTTACAAATAAAGCAATAGCATCACAAATTTCACAAATAAAGCATTTTT
TTCACTGCATTCTAGTTGTGGTTTGTCCAAACTCATCAATGTATCTTAACGCGTAAATTGTAA
GCGTTAATATTTTGTTAAAATTCGCGTTAAATTTTTGTTAAATCAGCTCATTTTTTAACCAATA
GGCCGAAATCGGCAAAATCCCTTATAAATCAAAAGAATAGACCGAGATAGGGTTGAGTGTT
GTTCCAGTTTGGAACAAGAGTCCACTATTAAAGAACGTGGACTCCAACGTCAAAGGGCGAA
AAACCGTCTATCAGGGCGATGGCCCACTACGTGAACCATCACCCTAATCAAGTTTTTTGGG
GTCGAGGTGCCGTAAAGCACTAAATCGGAACCCTAAAGGGAGCCCCCGATTTAGAGCTTG
ACGGGGAAAGCCGGCGAACGTGGCGAGAAAGGAAGGGAAGAAAGCGAAAGGAGCGGGC
GCTAGGGCGCTGGCAAGTGTAGCGGTCACGCTGCGCGTAACCACCACACCCGCCGCGCT
TAATGCGCCGCTACAGGGCGCGTCAGGTGGCACTTTTCGGGGAAATGTGCGCGGAACCC
CTATTTGTTTATTTTTCTAAATACATTCAAATATGTATCCGCTCATGAGACAATAACCCTGAT
AAATGCTTCAATAATATTGAAAAAGGAAGAGTCCTGAGGCGGAAAGAACCAGCTGTGGAAT
GTGTGTCAGTTAGGGTGTGGAAAGTCCCCAGGCTCCCCAGCAGGCAGAAGTATGCAAAGC
ATGCATCTCAATTAGTCAGCAACCAGGTGTGGAAAGTCCCCAGGCTCCCCAGCAGGCAGA
AGTATGCAAAGCATGCATCTCAATTAGTCAGCAACCATAGTCCCGCCCCTAACTCCGCCCA
TCCCGCCCCTAACTCCGCCCAGTTCCGCCCATTCTCCGCCCCATGGCTGACTAATTTTTTT
TATTTATGCAGAGGCCGAGGCCGCCTCGGCCTCTGAGCTATTCCAGAAGTAGTGAGGAGG
CTTTTTTGGAGGCCTAGGCTTTTGCAAAGATCGATCAAGAGACAGGATGAGGATCGTTTCG
CATGATTGAACAAGATGGATTGCACGCAGGTTCTCCGGCCGCTTGGGTGGAGAGGCTATT
CGGCTATGACTGGGCACAACAGACAATCGGCTGCTCTGATGCCGCCGTGTTCCGGCTGTC
AGCGCAGGGGCGCCCGGTTCTTTTTGTCAAGACCGACCTGTCCGGTGCCCTGAATGAACT
GCAAGACGAGGCAGCGCGGCTATCGTGGCTGGCCACGACGGGCGTTCCTTGCGCAGCTG
TGCTCGACGTTGTCACTGAAGCGGGAAGGGACTGGCTGCTATTGGGCGAAGTGCCGGGG
CAGGATCTCCTGTCATCTCACCTTGCTCCTGCCGAGAAAGTATCCATCATGGCTGATGCAA
TGCGGCGGCTGCATACGCTTGATCCGGCTACCTGCCCATTCGACCACCAAGCGAAACATC
GCATCGAGCGAGCACGTACTCGGATGGAAGCCGGTCTTGTCGATCAGGATGATCTGGACG
AAGAGCATCAGGGGCTCGCGCCAGCCGAACTGTTCGCCAGGCTCAAGGCGAGCATGCCC
GACGGCGAGGATCTCGTCGTGACCCATGGCGATGCCTGCTTGCCGAATATCATGGTGGAA
AATGGCCGCTTTTCTGGATTCATCGACTGTGGCCGGCTGGGTGTGGCGGACCGCTATCAG
GACATAGCGTTGGCTACCCGTGATATTGCTGAAGAGCTTGGCGGCGAATGGGCTGACCGC
TTCCTCGTGCTTTACGGTATCGCCGCTCCCGATTCGCAGCGCATCGCCTTCTATCGCCTTC
TTGACGAGTTCTTCTGAGCGGGACTCTGGGGTTCGAAATGACCGACCAAGCGACGCCCAA
CCTGCCATCACGAGATTTCGATTCCACCGCCGCCTTCTATGAAAGGTTGGGCTTCGGAATC
GTTTTCCGGGACGCCGGCTGGATGATCCTCCAGCGCGGGGATCTCATGCTGGAGTTCTTC

GCCCACCCTAGGGGGAGGCTAACTGAAACACGGAAGGAGACAATACCGGAAGGAACCCG
CGCTATGACGGCAATAAAAAGACAGAATAAAACGCACGGTGTTGGGTCGTTTGTTCATAAA
CGCGGGGTTCGGTCCCAGGGCTGGCACTCTGTCGATACCCCACCGAGACCCCATTGGGG
CCAATACGCCCGCGTTTCTTCCTTTTCCCCACCCCACCCCCAAGTTCGGGTGAAGGCCC
AGGGCTCGCAGCCAACGTCGGGGCGGCAGGCCCTGCCATAGCCTCAGGTTACTCATATAT
ACTTTAGATTGATTTAAAACTTCATTTTTAATTTAAAAGGATCTAGGTGAAGATCCTTTTTGAT
AATCTCATGACCAAAATCCCTTAACGTGA
